# Adrenergic and mesenchymal signatures are identifiable in cell-free DNA and correlate with metastatic disease burden in children with neuroblastoma

**DOI:** 10.1101/2023.08.30.554943

**Authors:** Omar R. Vayani, Maria E. Kaufman, Kelley Moore, Mohansrinivas Chennakesavalu, Rachel TerHaar, Gepoliano Chaves, Alexandre Chlenski, Chuan He, Susan L. Cohn, Mark A. Applebaum

**Affiliations:** The University of Chicago Pritzker School of Medicine, Chicago, IL, 60637, USA; Department of Pediatrics, Section of Hematology/Oncology, The University of Chicago, Chicago, IL, 60637, USA; Department of Chemistry, The University of Chicago, Chicago, IL, 60637, USA

**Keywords:** neuroblastoma, cell free DNA, epigenomics, mesenchymal, adrenergic

## Abstract

**Background:** Cell free DNA (cfDNA) profiles of 5-hydroxymethylcytosine (5-hmC), an epigenetic marker of open chromatin and active gene expression, are correlated with metastatic disease burden in patients with neuroblastoma. Neuroblastoma tumors are comprised of adrenergic (ADRN) and mesenchymal (MES) cells, and the relative abundance of each in tumor biopsies has prognostic implications. We hypothesized that ADRN and MES specific signatures could be quantified in cfDNA 5-hmC profiles and would augment the detection of metastatic burden in patients with neuroblastoma.

**Methods:** We previously performed an integrative analysis to identify ADRN and MES specific genes (n=373 and n=159, respectively). Purified DNA from cell lines was serial diluted with healthy donor cfDNA. Using Gene Set Variation Analysis (GSVA), ADRN and MES signatures were optimized. We then quantified signature scores, and our prior neuroblastoma signature, in cfDNA from 84 samples from 46 high-risk patients including 21 patients with serial samples.

**Results:** Samples from patients with higher metastatic burden had increased GSVA scores for both ADRN and MES gene signatures (p < 0.001). While ADRN and MES signature scores tracked together in serially collected samples, we identified instances of patients with increases in either MES or ADRN score at relapse.

**Conclusions:** While it is feasible to identify ADRN and MES signatures using 5-hmC profiles of cfDNA from neuroblastoma patients and correlate these signatures to metastatic burden, additional data are needed to determine the optimal strategies for clinical implementation. Prospective evaluation in larger cohorts is ongoing.

## INTRODUCTION

Neuroblastoma is the most common extracranial solid tumor of childhood. It originates from neural crest cells and most commonly presents in the adrenal glands.^1^ It is especially notable for its diverse clinical presentation, reflecting its biological heterogeneity. Currently, the Children’s Oncology Group risk stratification system uses a combination of clinical and biologic markers including International Neuroblastoma Risk Group stage, age, and *MYCN*-amplification status to determine optimal therapeutic approaches for patients.^2^ Those with unfavorable features including age over 18 months and *MYCN*-amplified tumors are categorized as high-risk and only 50% of these patients achieve long-term survival.^3^ Because high-risk neuroblastomas are clinically heterogenous, it is imperative to further understand their biologic nuances to promote the application of more refined therapies.^4^

High-risk neuroblastoma tumors consists primarily of adrenergic (ADRN) cells, which are rapidly dividing, neuronal in nature, and more easily eradicated with standard chemotherapy.^5^ Conversely, mesenchymal (MES) cells are thought to be more “stem-like,” slowing dividing, and less susceptible to chemotherapy.^5^ Each subtype has been shown to have lineage-specific chromatin landscapes and transcriptional programs. As these cellular identities are established by epigenetic regulation, it is possible for these chromatin marks to be modified, allowing for interconversion between MES and ADRN lineages.^6^ Furthermore, initial studies of ADRN and MES phenotypes identified increased MES cells in relapsed tumors.^5^ However, this biology is complex as MES signatures are also highly correlated with immune cell infiltration which may itself be a marker of improved outcomes.^7,8^

Among pediatric solid tumors, neuroblastoma patients have the highest levels of tumor derived cell-free DNA (cfDNA) in the blood, making this an ideal application of DNA based liquid biopsies to monitor disease and identify biologic variables that may influence response to therapy.^9,10^ Unlike tumor biopsies, liquid biopsies can be easily obtained serially with low cost and minimal safety concerns. Because the tumor material in blood disproportionately stems from the most aggressive metastatic cells, relapse driving genomic aberrations of tumors are enriched in peripheral blood.^11,12^ We previously demonstrated that 5-hydroxymethylcytosine (5-hmC) profiles can be readily obtained from cfDNA from children with neuroblastoma and are distinct from those from well children. 5-hmC is generated by the oxidation of 5-methylcytosine in DNA by the TET enzymes and is an epigenetic marker of open chromatin and active gene expression. The amount of 5-hmC deposition on a gene strongly correlates with RNA levels in neuroblastoma tumors.^13^ Furthermore, in a retrospective study 5-hmC profiles were correlated with metastatic disease burden in children and could be used to assess response to therapy, a well-known surrogate of outcome.^14^ Thus, we hypothesized that implementing quantitatively ADRN and MES specific signatures could be determined in cfDNA 5-hmC profiles, providing additional information about clincal disease burden and response while helping to stratify patients according to molecular remission status.

## METHODS

### 2.1. Patients and 5-hmC libraries from cfDNA data

We utilized data from our previously described cohort.^14^ In brief, all patients were diagnosed with neuroblastoma between the ages of 0 and 29 years and samples were collected at two local children’s hospitals after approval by local Institutional Review Boards. Response to treatment was assessed according to the 2017 International Neuroblastoma Response Criteria (INRC).^15^ Patients were excluded if they did not have high-risk disease. Blood samples were collected in EDTA tubes and processed within two hours of collection. FastQ files were obtained from 5-hmC library preparation and sequencing as described. Data are available in dbGaP (phs001831.v1.p1). We previously performed unsupervised hierarchical clustering of 5-hmC profiles to identify four clusters of samples, correlating with metastatic burden.^14^ These four clusters were enriched for groups of patients who had either high, medium, low, or no clinically active metastatic disease according to cfDNA 5-hmC profiles.

### 2.2. Culture of neuroblastoma cell lines

Five ADRN neuroblastoma cell lines (LA1-55n, SH-SY5Y, NBL-W-N, SK-N-BE2, and NBL-S) and three MES lines (LA1-5s, SHEP, and NBL-W-S) were grown at 5% CO2 in RPMI1640 medium (Life Technologies) supplemented with 10% heat-inactivated FBS, 2 mmol/L l-glutamine, and 1% penicillin/streptomycin. Non-adherent ADRN cell lines with the potential of phenotype switching were disassociated from the flask surface using a short tapping technique and collected from solution for subculturing. For adherent MES cell lines, the media was removed after short tapping and cells were subsequently dissociated from flasks by trypsinization. To ensure the phenotype lineage of the cell lines, cells were observed for phenotypic switching throughout.

### 2.3. Nano-hmC-Seal library preparation and sequencing

Cells were lysed and DNA was obtained using Qiagen Purgene® Core Kit A. To replicate clinically heterogenous cfDNA which is derived from an admixture of MES and ADRN cells, 100ng of DNA from each cell line was combined and an aliquot of 150ng was fragmented using KAPA Frag Enzyme (Kapa Biosystems KK8514) and KAPA Frag Buffer (10x) (Kapa Biosystems KK8514) at 4°C for 30 minutes. Fragmented DNA was purified using ZYMO DNA Clean & Concentrator™-5 kit and quantified using Qubit fluorometer (Life Technologies). Purified DNA was serial diluted with healthy donor cfDNA to generate libraries from 100%, 50%, 25%, 12.5%, 6.25%, 3.13%, 1.56%, and 0% neuroblastoma DNA, respectively. Nano-hmC-Seal libraries were constructed from 10 ng of DNA for each dilution in duplicate and sequenced as described.^16,17^

### 2.4 5-hmC data processing

For the 5-hmC cfDNA data, FASTQC v0.11.5 was used to examine sequence quality.^18^ Raw reads were processed using Trimmomatic and aligned to hg38 using Bowtie2 v.2.3.0 using default settings.^19,20^ Duplicate reads were marked and removed with Picard v2.8.1.^21^ Aligned reads with Mapping Quality Score >= 10 were counted using featureCounts.^22^

### 2.5. Determination of cell lineage specific genes

We previously generated chromatin immunoprecipitation (ChIP) data from the same eight cell lines for H3K27Ac and H3K4me3 with matched RNAseq, and KAS-seq as described.^23^ To identify cell lineage specific genes, we first compiled ADRN and MES-specific super enhancers using Rank Ordering of Super Enhancers (ROSE) on H3K27Ac and H3K4me3 profiles.^24,25^ We then used the KAS-seq data and pipeline to determine which super-enhancers were single-stranded.^26^ Genes differentially expressed between MES and ADRN cell lines were identified (p_adj_ < 0.05 and absolute log fold change > 1) using DESeq2 v1.20.0 in R v3.6.0.^27^ Lineage specific differentially expressed genes within 500 kb of a single stranded super enhancers (ssSEs) were then selected for each signature. A comprehensive evaluation and validation of these signatures in multiple cohorts of diagnostic neuroblastoma biopsies has been reported.^23^

### 2.6. Identifying and optimizing ADRN and MES signatures in cell line 5-hmC profiles

Read counts for the entire gene body were loaded into DESeq2.^27^ Gene Set Variation Analysis (GSVA),^28^ a non-parametric, unsupervised gene set enrichment analysis method, was implemented using default arguments for each sample and plotted against percent neuroblastoma DNA. Pearson correlations between neuroblastoma DNA concentration and GSVA score were determined. To optimize the correlation of MES and ADRN GSVA scores with percent tumor DNA and remove genes with elevated 5-hmC deposition in normal blood, we performed a leave one out analysis in which individual genes were removed and a new correlation slope was generated. If the correlation was lower after removal of a gene, that gene was kept. If the correlation was improved after removing a gene, it was dropped. The optimized ADRN and MES signatures and a non-lineage specific signature previously identified was then assessed in patient samples using GSVA.^23^ Pathway analysis and compilation were completed using Enrichr.^29–31^ GSVA computed scores were compared across groups using the Kruskal-Wallis test.

### 2.7. ROC analysis and associations with 5-hmC profiles and disease progression

A Pearson correlation was used to assess the relationship of GSVA scores for each signature. ROC analysis and area under the curve (AUC) calculation for the each signature was completed using plotROC and pROC packages in R. DeLong’s test was used to assess differences in the curves and their AUCs.^32–34^

## RESULTS

### 3.1. MES and ADRN signatures in serial dilutions

The initial gene signatures included 373 ADRN specific genes and 159 MES specific genes. The linear correlation of GSVA scores of these signatures with increasing percent tumor DNA was 0.93 and 0.41, respectively (**Figure 1A and 1B**). MES cells are enriched for immune signatures,^8^ and we examined the effect of overlap of genes in our ADRN and MES signatures with those from normal peripheral blood which is enriched for immune signatures.^12^ Using a leave one out analysis, 156 genes and 84 genes were removed from the ADRN and MES gene lists, respectively (**Supplementary Tables 1 and 2**). The removed ADRN genes were enriched for pathways including organelle and cytoskeletal organization (**Supplementary Table 3**) whereas the removed MES genes were enriched for pathways of endothelium and immune cells (**Supplementary Table 4**). Top pathways for the remaining 217 ADRN genes included expected pathways of neuronal function (**Supplementary Table 5**), whereas the 75 remaining MES genes were enriched for decreased cell proliferation and increased DNA transcription (**Supplementary Table 6).** The final correlation between ADRN and MES signatures and increasing percent tumor DNA with the remaining genes was 0.98 and 0.99, respectively (**Figure 1B and 1D**). We compared our gene sets to those previously published by van Groningen et. al..^5^ 33 ADRN genes (representation factor = 8.2, p < 7.721e-21) and 9 MES genes (representation factor = 4.9, p < 8.426e-05) overlapped, suggesting significant similarities between ours and others MES and ADRN signatures.

**Figure 1.**
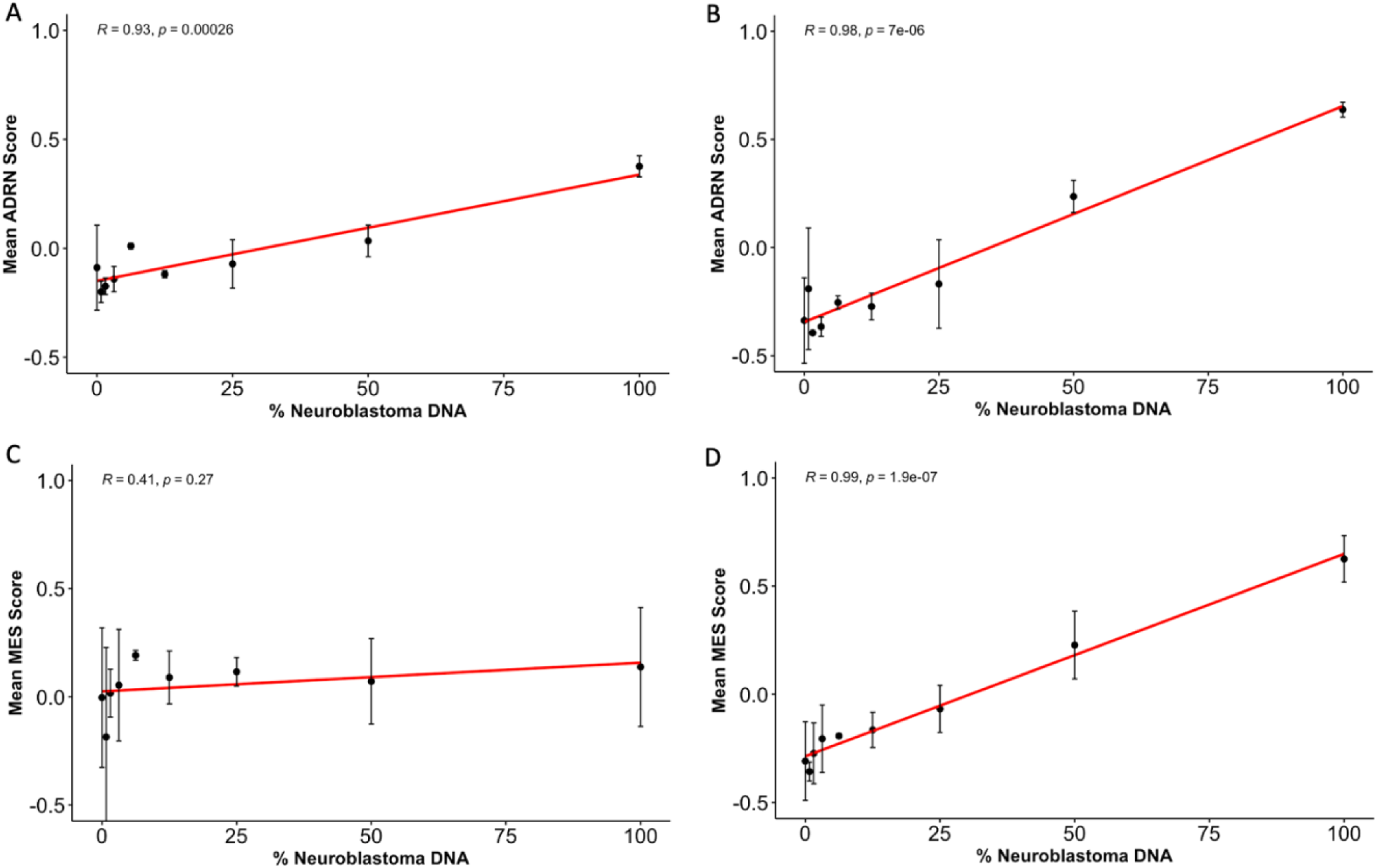
ADRN and MES scores by neuroblastoma DNA percent concentration. **A)** ADRN score by neuroblastoma DNA percent concentration prior to the leave one out analysis (R^2^ = 0.93); 373 genes. **B)** Optimized ADRN score (R^2^ = 0.98); 217 genes. **B)** MES score by neuroblastoma DNA percent concentration prior to the leave one out analysis (R^2^ = 0.41); 159 genes. **D)** Optimized MES score (R^2^ = 0.99); 75 genes.

We also utilized a previously generated signature consisting of 450 genes that had higher 5-hmC deposition on genes from samples with high metastatic disease burden compared to those with no metastatic disease burden seen on clinical imaging or bone marrow biopsies.^13^ We defined this as the NBL signature (**Supplementary Table 7**). Of the 450 genes in the NBL signature, 58 (12.8%) also had increased expression in ADRN cell lines compared to MES cell lines whereas 47 (10.4%) had increased expression in MES cell lines compared to ADRN cell lines. 345 genes (76.8%) were not significantly differentially expressed between ADRN and MES cells. Despite having low numbers of overlapping ADRN genes, the 450 genes were enriched for neuronal pathways commonly observed in this phenotype (**Supplementary Table 8**).

### 3.2. MES and ADRN signatures in patient cfDNA

We utilized 84 cfDNA samples from 46 children with high-risk neuroblastoma collected between August 2016 and July 2019. Of the samples, 12 were categorized as high metastatic disease burden at the time of collection; 9 samples were categorized as moderate metastatic disease burden; 12 samples were categorized as low metastatic disease burden; and 51 samples were categorized as no metastatic disease burden as previously defined.^14^ 21 patients had 2 or more cfDNA samples collected at different time points.

Although the ADRN, MES, and NBL signatures were comprised of divergent genes, GSVA scores for all three signatures were significantly correlated. The correlation of the NBL signatures with ADRN signatures was 0.66 (p < 0.001, **Figure 2A**), while correlation for the ADRN signatures with MES signatures was 0.7 (p < 0.001, **Figure 2B**). GSVA scores of the NBL signatures and MES signatures had a correlation coefficient of 0.46 (p < 0.001, **Figure 2C**). An additional combined score was also evaluated using the established ADRN and MES scores. As these scores were strongly correlated to ADRN signature scores (r=0.99; **Supplementary** Figure 1), it was not further evaluated. Samples from patients categorized as having higher metastatic burden had increased GSVA enrichment scores regardless of the gene signature used (p < 0.001, **Figure 3**). However, higher enrichment scores were seen in samples from patients categorized with high disease burden using both the mean NBL (0.53) and the mean ADRN (0.52) signatures compared to the mean MES signature enrichment score (0.23, p < 0.001 and p < 0.001, respectively). Conversely, in samples from patients categorized as no disease burden, mean MES enrichment scores were lower compared to mean NBL enrichment scores (−0.14 versus −0.09, p = 0.017), but higher than mean ADRN enrichment scores (−0.14 versus −0.27, p < 0.001). 5-hmC deposition of representative ADRN (HAND1 and ISL1) and MES (EGFR and COL1A1) are shown in **Supplementary** Figure 2. These patterns suggest that both the level of disease burden and the balance of ADRN and MES cellular states may influence gene signature correlations. It appears that a higher disease burden tends to increase the correlation between NBL and ADRN signatures more than MES, whereas the absence of disease burden reveals a stronger correlation with ADRN than MES signatures. These observations could imply nuanced roles of ADRN and MES states across different stages of metastatic burden, which we further examined.

**Figure 2.**
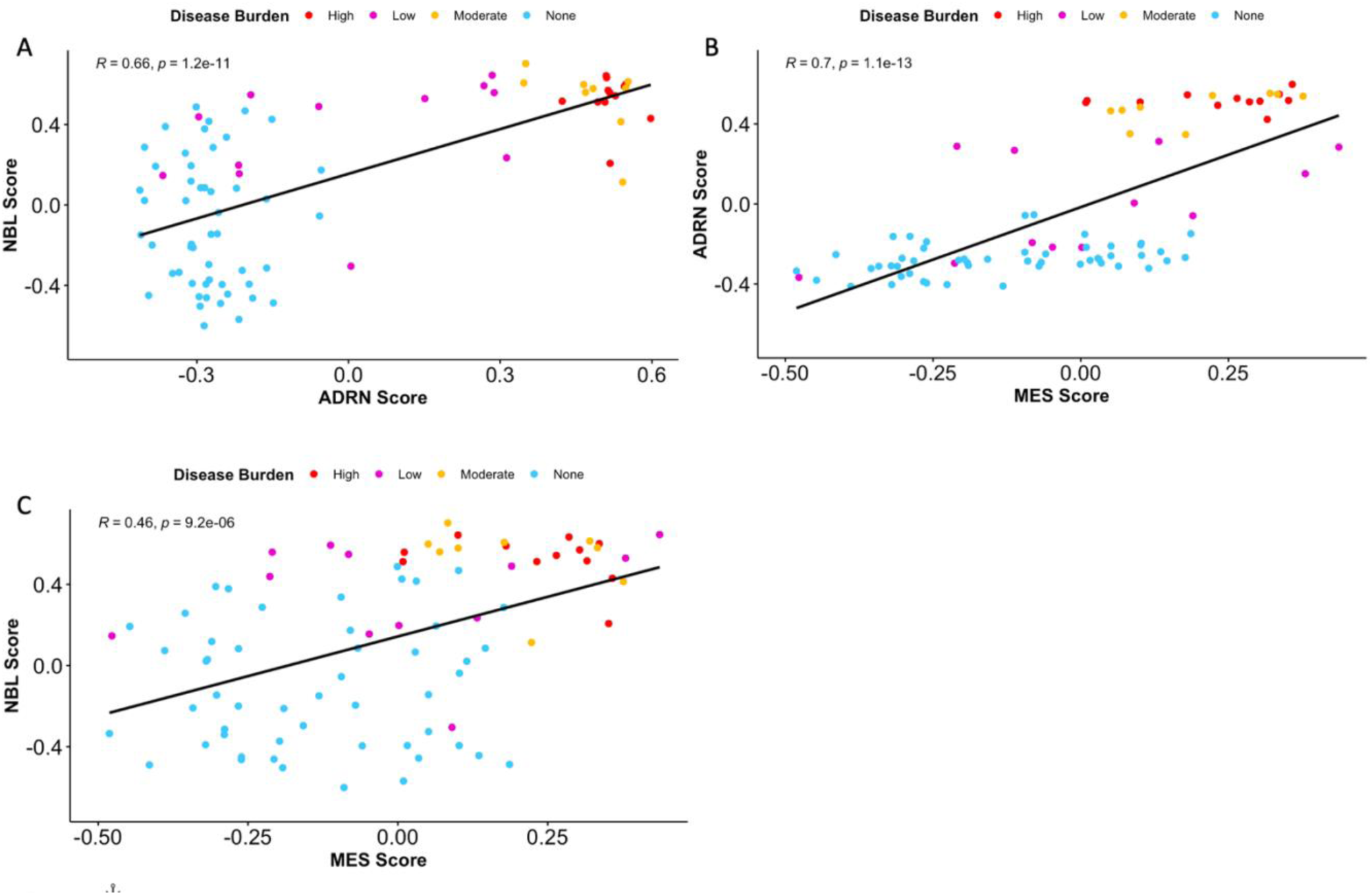
GSVA scores of the three gene signature sets are highly correlated across samples. The ADRN signature was highly correlated with both the NBL (A) and the MES (B) signature. Weaker but significant correlation was seen between the NBL and MES signatures.

**Figure 3.**
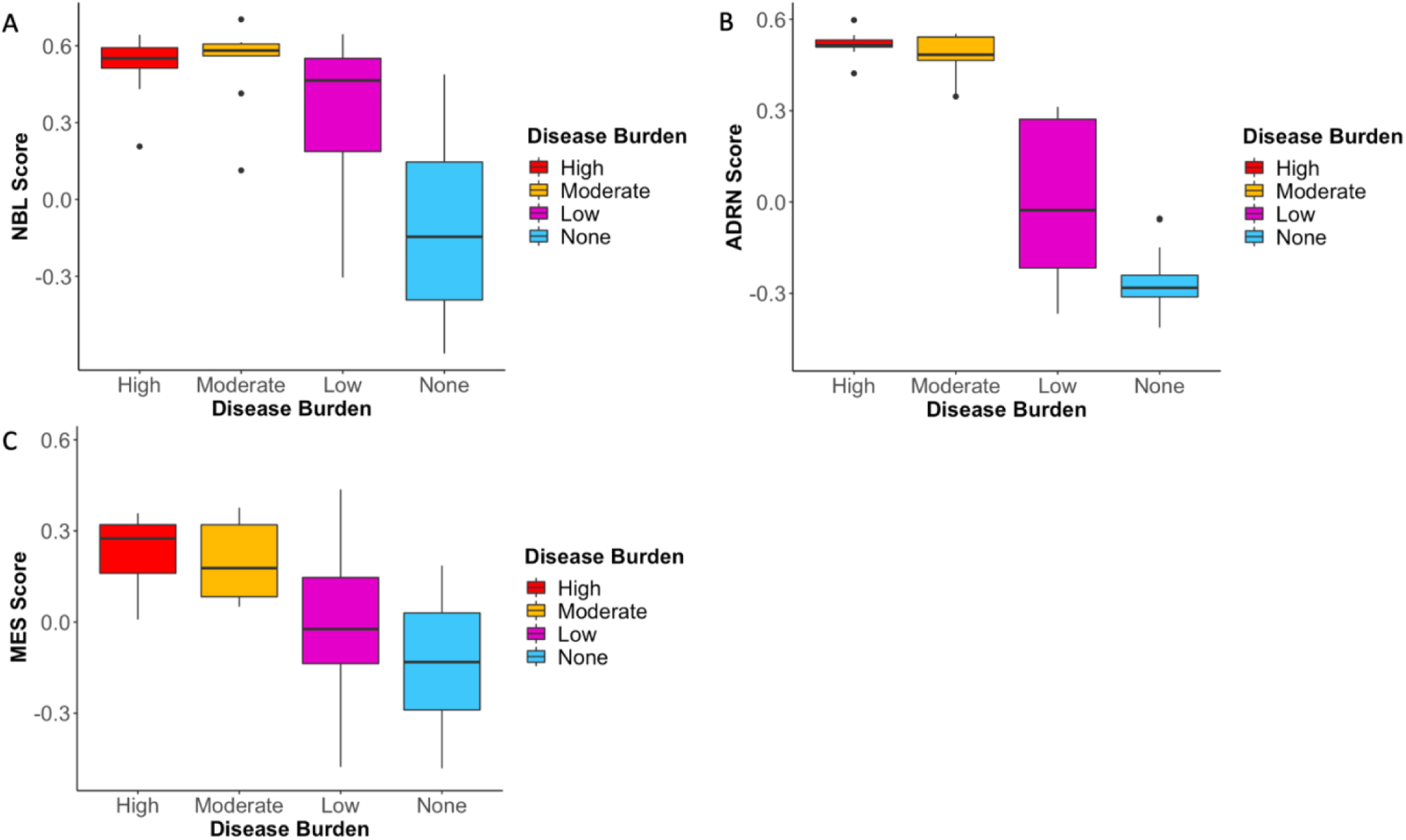
Mean signature score by neuroblastoma disease burden. NBL (A), ADRN (B), and MES (C) signature scores were highest in patients categorized with the largest burden of disease.

### 3.3. Association of signatures and subsequent disease progression

We next performed a ROC analysis to determine the potential utility for each scoring system to identify patients likely to experience disease progression (**Figure 4**). The area under the curve (AUC) for the NBL signature compared to ADRN signature was not significantly different (0.76 versus 0.72, p = 0.46). While both the NBL and ADRN signatures had higher AUC than the MES signature (AUC = 0.67), the difference was not significant (p = 0.22 and p = 0.14, respectively). The non-significant differences observed between the AUC values of three signatures is likely attributed to the high correlation of the underlying signatures to each other and that many of the samples were collected months before relapse. However, the AUC values are moderate to high, indicating potential clinical utility in disease progression identification.

**Figure 4.**
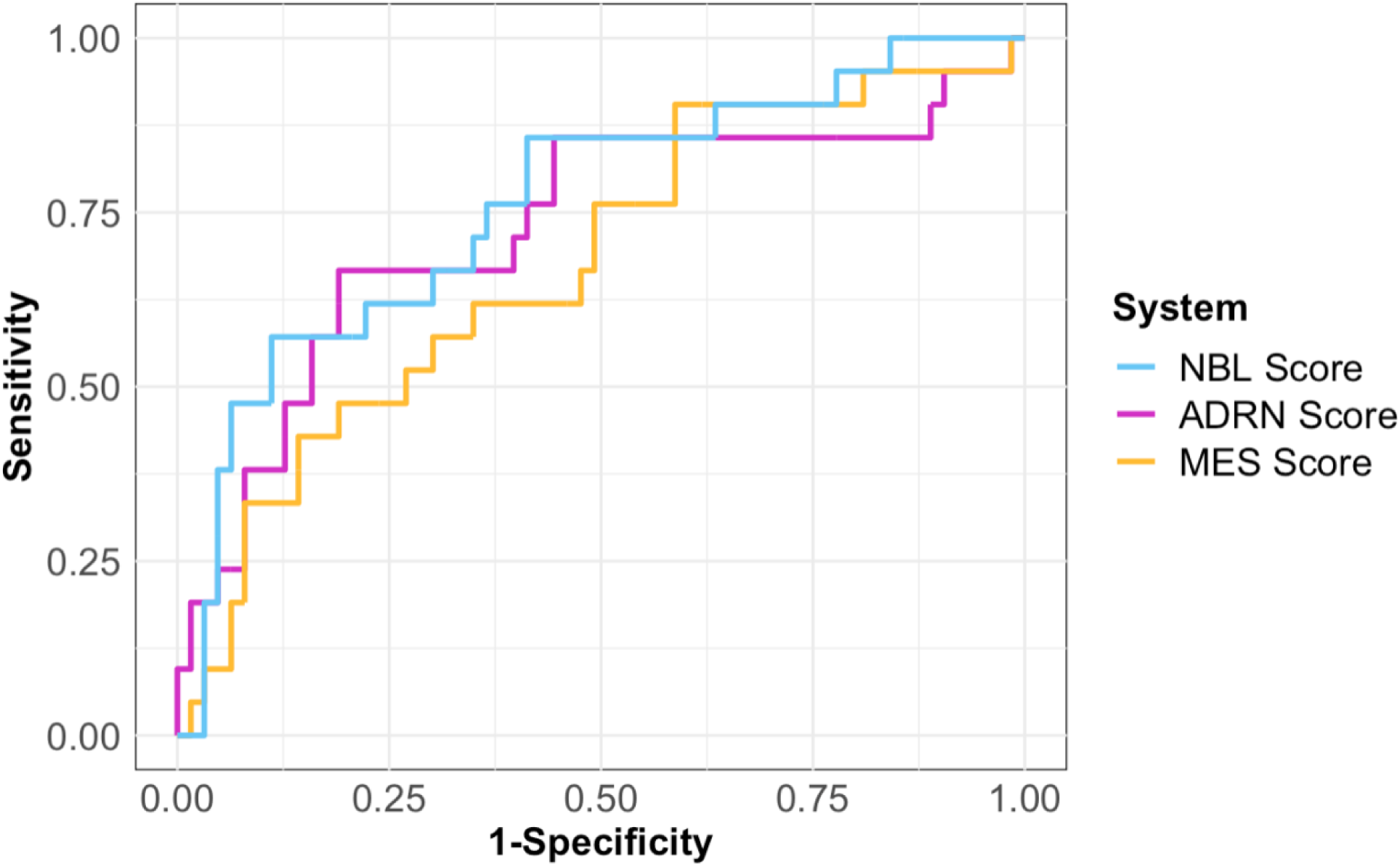
ROC analysis of the NBL, ADRN, and MES scoring systems. Area under the curve for each scoring system is 0.7626, 0.7263, and 0.6749, respectively.

### 3.4. Serial ADRN and MES associate with clinical response to treatment in patients with neuroblastoma

We previously established that hierarchical clustering using the NBL signature could be used to identify patients according to therapeutic response.^14^ To evaluate if GSVA scores using ADRN and MES signatures provide a more precise measure of response, we evaluated signature scores in 21 high-risk patients with more than 2 samples obtained at different time points (**Figure 5**). In 14 patients, both ADRN and MES signature scores tracked closely together regardless of outcome. Of the seven patients in which the scores did not track closely, only the MES score increased at relapse in one patient and only the ADRN score increase in one patient with progressive disease. In the other five patients, the MES scores increased in patients who remained in remission while the ADRN scores remained stable or decreased.

**Figure 5.**
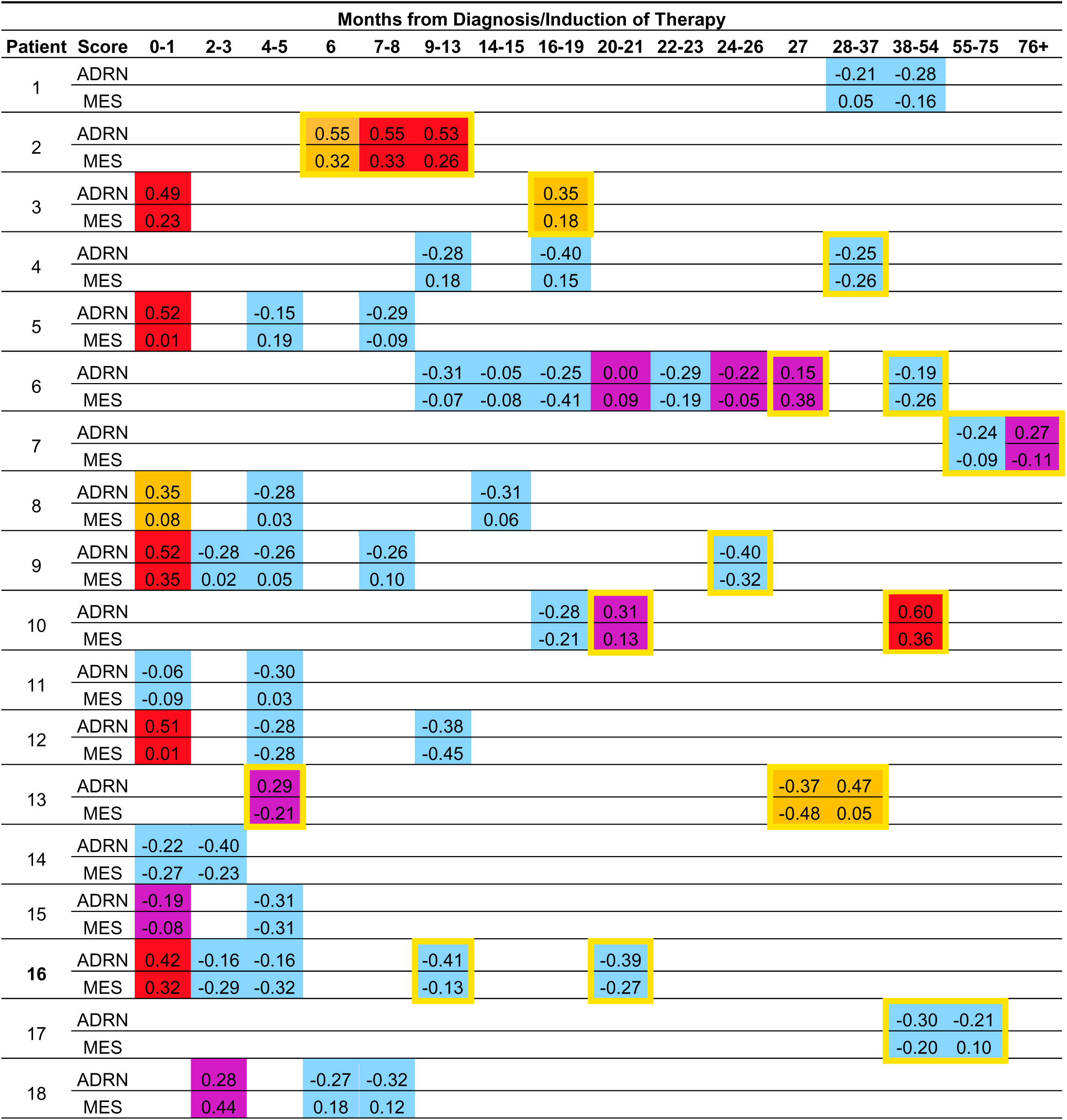
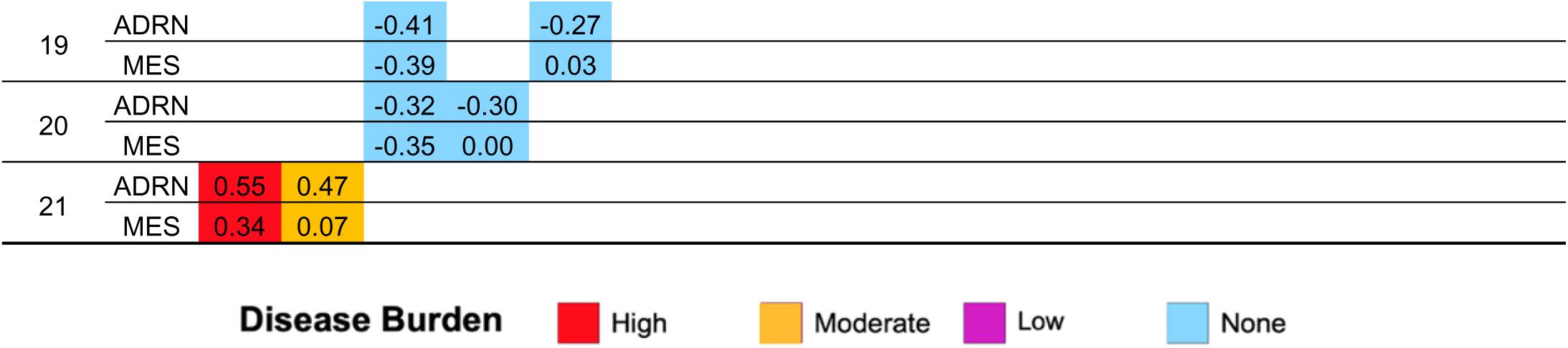
Individual MES and ADRN scores correlated with disease burden over time in 23 patient serial samples. Yellow boxes denote that the patient sample was taken after patient had relapsed. Patient 16 was a unique case in which the MES score was superior to ADRN scores at identifying recurrent disease.

## DISCUSSION

This study represents an exploration into the utility of ADRN and MES lineage-specific signatures in the context of cfDNA 5-hmC profiles for neuroblastoma patient monitoring. We generated novel ADRN and MES lineage specific gene signatures for use in cfDNA that were derived by incorporating differential expressed genes near super-enhancers and single stranded combined with transcriptionally engaged genes. These signatures can be detected in 5hmC cfDNA profiles generated from neuroblastoma patient blood samples, and we found that ADRN and MES scores were correlated with metastatic disease burden. As ADRN and MES scores were highly correlated (r = 0.7), they largely followed similar trends as patients went through treatment for primary or relapsed disease. However, we identified a small number of patients in which either the ADRN or the MES signature was associated with the presence or absence of metastatic disease burden. These results demonstrate that integrating lineage specific markers into a scoring system using 5-hmC can provide quantitative data that is informative of tumor burden and clinical relapse in patients. While we have previously shown that hierarchical clustering using a general neuroblastoma gene signature can segregate cfDNA samples according to metastatic disease burden, this study shows that quantitative single sample enrichment scoring can assess lineage specific circulating neuroblastoma DNA enabling the development of liquid biopsy-based biomarkers for this disease.

Biologic heterogeneity is a hallmark characteristic of neuroblastoma. Importantly, neuroblastoma cells are capable of interconverting between MES and ADRN with corresponding changes in phenotype and chemosensitivity.^5,35,36^ While the precise association between heterogeneity and therapeutic response in newly diagnosed patients is yet to be determined, recent studies demonstrated that neuroblastomas enriched with MES cells associated with relapse, and several studies have focused on converting cellular phenotype to alter therapeutic sensitivity.^37–40^ However, there are also now multiple reports suggesting that increased MES signature scores in diagnostic biopsies are associated with improved patient outcomes, suggesting the clinical ramifications of these phenotypes are not clear cut.^37,41^ In our patient cohort, 14 of 21 patients had ADRN and MES scores that tracked together, regardless of disease status. Of the seven patients where MES and ADRN scores diverged over time, two were associated with relapse, suggesting that there may be patients for whom differing lineages predominate at relapse. However, we also noted rising MES scores in four patients who remained in remission, suggesting the need for further optimization of this signature.

Should the findings in this study be validated in prospective studies, the observed divergence between ADRN and MES signatures might represent adaptive changes within the tumor microenvironment in response to therapy, where certain lineages might be favored under certain conditions depending on the tumor characteristics and microenvironment.^42,43^ However, neuroblastoma tumors during diagnosis or relapse are heterogenous and comprised of both ADRN and MES cells,^5^ and this may contribute to our observation that MES and ADRN signatures track closely in all patients in most of the samples analyzed. Improvements in our understanding of neuroblastoma evolution over the course of treatment and relapse will likely enable more precise deployments of lineage specific signatures in the optimal context.

While others have identified lineage specific gene signatures, in addition to evaluating genes near super-enhancers with differential expression in ADRN vs MES cells, we also utilized KAS-seq data and incorporated genes with single stranded enhancers, a marker of active transcription, that were differential expressed in the phenotypically distinct neuroblastoma cell types.^5,26,44^ By optimizing these signatures using serial dilutions of neuroblastoma DNA with healthy donor cfDNA we have demonstrated the feasibility of detecting low levels of neuroblastoma-derived cfDNA in patients with neuroblastoma. Our results demonstrate that the ADRN signature had a wider distribution of enrichment scores across the spectrum of metastatic disease burden but similar kinetics over time was observed with MES signatures. These findings are consistent with mRNA based MES and ADRN signatures in peripheral blood and bone marrow from children with neuroblastoma.^45–47^ The highest levels of the ADRN mRNA signature were detected at diagnosis and relapse in the cohort described by van Wezel, and decreased expression was observed during treatment. In contrast, the MES mRNA signature increased during treatment and was associated with relapse in one patient. These investigators detected the MES mRNA signature in only 14 out of 27 patients at diagnosis and rarely at the time of relapse. As circulating cfDNA is protected from degradation by nucleosomes, our signatures may be improved in the future by using nucleosome capture of DNA from cell lines prior to sequencing.^48^ Furthermore, combining this approach with other methods such as ultra-low passage whole genome sequencing, mRNA, or methylation signatures may provide synergistic sensitivity to detect minimal residual disease.^10,47,49,50^

Although this study is limited by a lack of validation cohort, we have previously shown that 5-hmC profiles are consistent and robust across both discovery and validation cohorts.^14^ We also previously demonstrated that 5-hmC profiles can identify cell pathways that the biology of neuroblastoma. We now demonstrate markers associated with phenotypically distinct neuroblastoma cell states can be used to further categorize neuroblastoma cfDNA 5-hmC profiles. While the AUC for detection of subsequent relapse was suggestive of the prognostic ability of ADRN and MES signatures, further refinement is needed. Another consideration in our approach was the use of cultured cell lines. We recognize the potential for primary patient-derived cell lines or fresh tissue samples to be of higher fidelity to *in situ* disease. Improved processing of genomic DNA from cell lines to more closely mimic cfDNA using nucleosome and ADRN and MES specific serial dilutions combined with machine learning approaches will likely be able to further improve the prognostic value of these gene signatures.

Future studies will focus on prospectively collecting patient samples at standardized timepoints to further validate and evaluate the clinical utility our findings. These studies will aim to delineate which epigenetic landscapes are driver adaptations versus passengers in the progression and relapse of children with neuroblastoma. Through them, we aim to define an informative and clinically applicable signature for patient management.

## CONCLUSIONS

We show that it is feasible to identify ADRN and MES signatures in cfDNA 5-hmC profiles from patients with neuroblastoma. If validated, these signatures may identify clinical scenarios in which they are better able to detect progressive disease. Prospective evaluation of this approach is ongoing as part of a Children’s Oncology Group phase III trial.

## Author Contributions

Omar R. Vayani: formal analysis, writing-original draft, writing-review and editing. Maria Kaufman: writing-review and editing. Kelley Moore: data curation and writing-review and editing. Mohansrinivas Chennakesavalu: formal analysis and writing-review and editing. Gepoliano Chaves: writing-review and editing. Alexandre Chlenski: data curation and writing-review and editing. Chuan He: writing-review and editing. Susan L. Cohn: writing-review and editing and supervision. Mark A. Applebaum: conceptualization, data curation, writing-original draft and writing-review and editing, and supervision.

## Funding

This work was supported in part by the Alex’s Lemonade Stand Foundation (MAA); a kind gift from Barry & Kimberly Fields (SLC and CH); the Matthew Bittker Foundation (SLC); a gift from the MacRitchie Family (SLC), and the National Institutes of Health, R37CA262781 (MAA) and R25CA240134 (ORV).

## Institutional Review Board Statement

The study was approved by the local Institutional Review Board according to the U.S. Common Rule ethical guidelines.

## Informed Consent Statement

Written informed consent was obtained from parents or guardians.

## Data Availability Statement

Data are available in dbGaP (phs001831.v1.p1)

## Conflicts of Interest

Susan L. Cohn: stock ownership in Pfizer, Merck, and Lilly; served on an advisory board for YMabs Therapeutics. Mark A. Applebaum: consultancy fees from Illumina Radiopharmaceuticals. The remaining authors made no disclosures.

## Abbreviations

cfDNA: Cell-free DNA
ADRN: Adrenergic
MES: Mesenchymal
ssSE: Single stranded super enhancer
GSVA: Gene Set Variation Analysis
AUC: Area under the curve

## Notes

https://www.ncbi.nlm.nih.gov/projects/gap/cgi-bin/study.cgi?study_id=phs001831.v1.p1

