## Supplementary Figures 1 and 2 for "Adrenergic and mesenchymal signatures are identifiable in cell-free DNA and correlate with metastatic disease burden in children with neuroblastoma"

**Supplementary Figure 1. GSVA scores of the Combined signature is correlated with the ADRN and MES signatures.** (A) The ADRN signature was highly correlated with the Combined signature. (B) Weaker but significant correlation was seen with the MES signature.

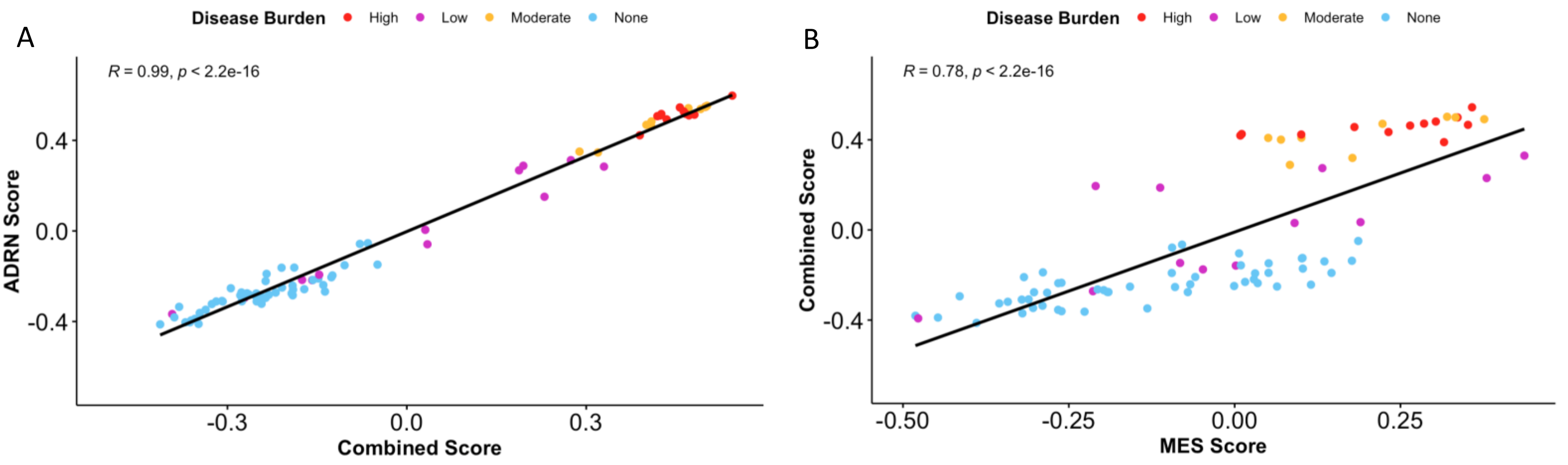

**Supplementary Figure 2. Mean number of reads by neuroblastoma disease burden.** (A) HAND1 and (B) ISL1 are ADRN signature genes while (C) EGFR and (D) COL1A1 are MES signature genes.

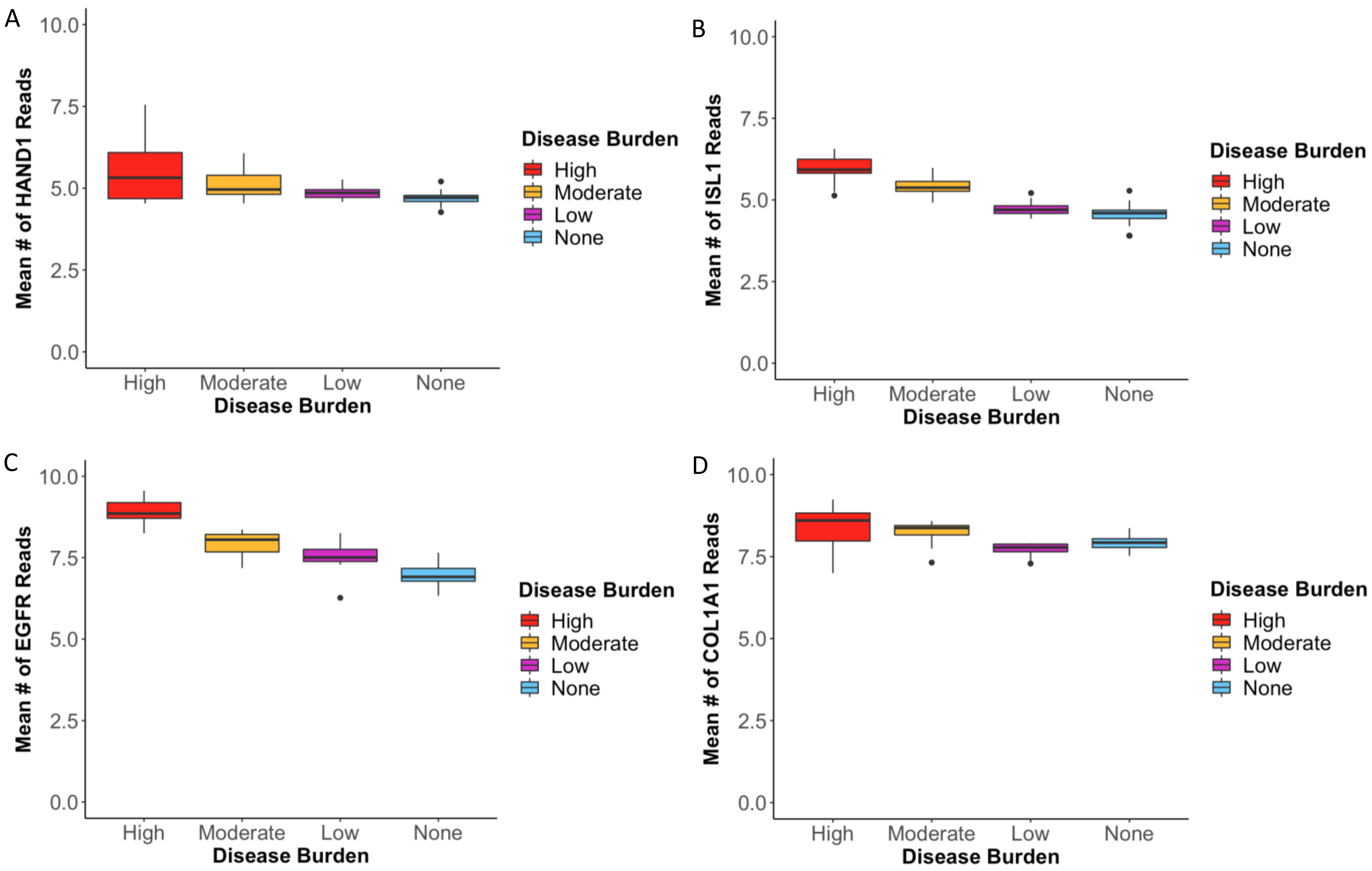
